# CINS: Cell Interaction Network inference from Single cell expression data

**DOI:** 10.1101/2021.02.22.432206

**Authors:** Ye Yuan, Carlos Cosme, Taylor Sterling Adams, Jonas Schupp, Koji Sakamoto, Nikos Xylourgidis, Matthew Ruffalo, Naftali Kaminski, Ziv Bar-Joseph

## Abstract

Studies comparing single cell RNA-Seq (scRNA-Seq) data between conditions mainly focus on differences in the proportion of cell types or on differentially expressed genes. In many cases these differences are driven by changes in cell interactions which are challenging to infer without spatial information. To determine cell-cell interactions that differ between conditions we developed the Cell Interaction Network Inference (CINS) pipeline. CINS combines Bayesian network analysis with regression-based modeling to identify differential cell type interactions and the proteins that underlie them. We tested CINS on a disease case control and on an aging human dataset. In both cases CINS correctly identifies cell type interactions and the ligands involved in these interactions. We performed additional mouse aging scRNA-Seq experiments which further support the interactions identified by CINS.

## Introduction

The ability to profile the expression of genes at the single cell level has revolutionized gene expression studies. Single cell RNA-Seq (scRNA-Seq) studies resulted in insights related to the cell type composition of tissues (1, 2), changes in cell type composition in various diseases and states (3), various differentiation pathways used within cells (4) and more. However, while scRNA-Seq provides valuable information about expression within cells, it is hard to use it to study interaction between cells. The main problem is that once cells are extracted it is very challenging to determine the spatial relationships among them (5).

Recently, a number of technologies have emerged for profiling single cell expression data with spatial resolution (6–9). These technologies often combine Fluorescence in situ hybridization (FISH) techniques with rapid sequencing technologies to provide information on the spatial expression of thousands of genes at various resolutions (10, 11). A number of recent computational methods have been developed to allow for the study of signaling pathways involved in cell-cell interactions from this type of spatially-resolved expression data (12, 13). However, while spatial transcriptomics studies are promising there are several challenges involved in employing this technique to study intercellular interactions. First, these techniques are still in their infancy and most labs do not have access or ability to perform such studies at the single cell resolution. More importantly, spatial transcriptomics often requires the fixation of the samples which limits their usage and can negatively impact their ability to accurately profile molecular quantities (10). In addition, spatial transcriptomics methods can scan only a small region of the tissue and so cannot be applied to large number of conditions and samples that are studied using scRNA-Seq.

Here we present a new method, the Cell Interaction Network Inference (CINS) pipeline, that enables researchers to study cell type interactions in scRNA-Seq data. CINS involves two major steps. First, it uses scRNA-Seq data from multiple samples of a similar condition (i.e. disease, age, etc.) to learn Bayesian networks which highlight the cell types whose distributions are co-varying under different conditions. Next, for the significant interactions identified in the Bayesian network analysis, CINS learns a regression model with ligand-target interaction matrix (14) that identifies the key ligands and targets that participate in the interactions between these cell types.

We tested CINS by applying it to both, disease and aging datasets. Using results of prior studies we show that CINS correctly identifies known interacting cell type pairs and ligands associated with these interactions. We also discuss several novel predictions made by CINS. Finally, we show that a number of CINS predicted cell type interactions are supported by a new scRNA-Seq lung aging dataset we profiled.

## Results

### The Cell Interaction Network Inference (CINS) Pipeline

We developed the Cell Interaction Network Inference (CINS) pipeline which uses single cell (sc) RNA-seq expression data to infer cell-cell interactions (**Fig. 1**). Given repeated experiments of the same condition / system CINS uses annotated cell type information to construct a Bayesian network (BN) that models causal relationships between different cell types. For this, CINS first discretizes the proportion data for each cell type using a Gaussian Mixture Model (GMM) with only two components and then learns a BN that models the joint probability distribution of the cell type mixtures observed for each sample. Significant causal relationships are determined based on bootstrapping. Next, for each of the significant pairs identified we infer the molecular pathways involved in the interactions by learning a ligand-target regression (LTR) model with ligand-target interaction database from NicheNet (14). The LTR model aims to explain changes in target genes as a function of changes in their activating ligands allowing CINS to identify the most significant ligands that regulate the cell-cell interactions.

**Figure 1.**
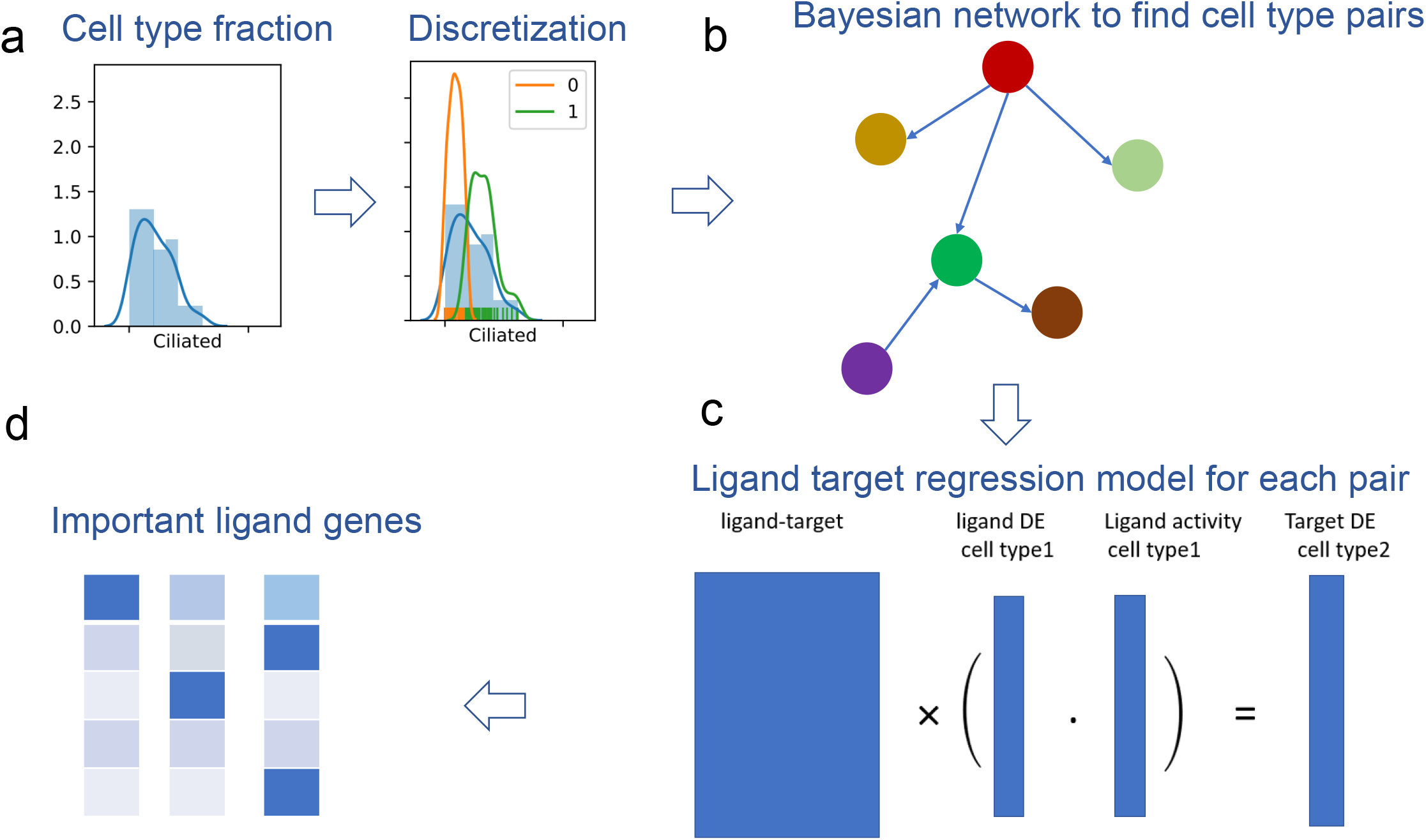
Overview of CINS. (a) Cell type information is discretized by learning Gaussian Mixture Model (GMM) for the fraction of cells of this type in each sample. (b) A Bayesian network (BN) is learned using the discretized cell abundance information. Bootstrapping is performed to identify significant interactions between cell types. (c) For pairs identified in the directed bootstrap BN analysis, a ligand-target regression (LTR) model is learned. In this model we use expression of ligands in the cell type with the outgoing edge to predict the expression of targets genes in the cell type with incoming edge. (d) Finally, LTR is used to select key ligands that underlie the cell-cell interactions identified in the BN. cell interaction.

### Inferring cell type interactions using Bootstrapped Bayesian Network

We first studied a lung disease scRNA-Seq dataset (15). The lung disease dataset contained scRNA-Seq data for 28 healthy (controls) and 32 Idiopathic Pulmonary Fibrosis (IPF) individuals. A total of 250,942 cells were profiled for these individuals. Cell type annotations were assigned based on the original study and we used the detailed assignments that provided information on 39 cell types.

We used CINS to explore differential cell type interactions between IPF and control samples. For this, we constructed two different networks based on the cells profiled for each condition. We next performed bootstrap analysis to determine the significance of each edge in each condition. Edges that appear in the majority of bootstrap iterations likely represent real relationships in the data rather than noise (16, 17). Resulting BNs for the two conditions are presented in **Fig. 2A&B.** As the figures show, there are some edges that appear for both conditions. These include Basal to Goblet cell interactions, which agrees with the fact that club cell’s attachment sites are provided by Basal cell (18). However, there are also many differences between edges selected for the two condition networks. **Tab. 1** summarized the top differences based on the signed difference in edge count in 100 bootstrap iterations for IPF and control (See **Tab. S1** for differences for all detected edges). Several of the highest scoring edges are supported by prior work. For example, the edge from Treg to Fibroblast cell is supported by a previous study suggesting that Treg’s can negatively regulate fibroblast activity (19). The edge between cDC2 and cDC1 is also supported by recent work showing that cDC2 and cDC1 are cross-talking with each other (20). Several other top scoring edges are supported by the literature as referenced in **Tab. 1.**

**Figure 2.**
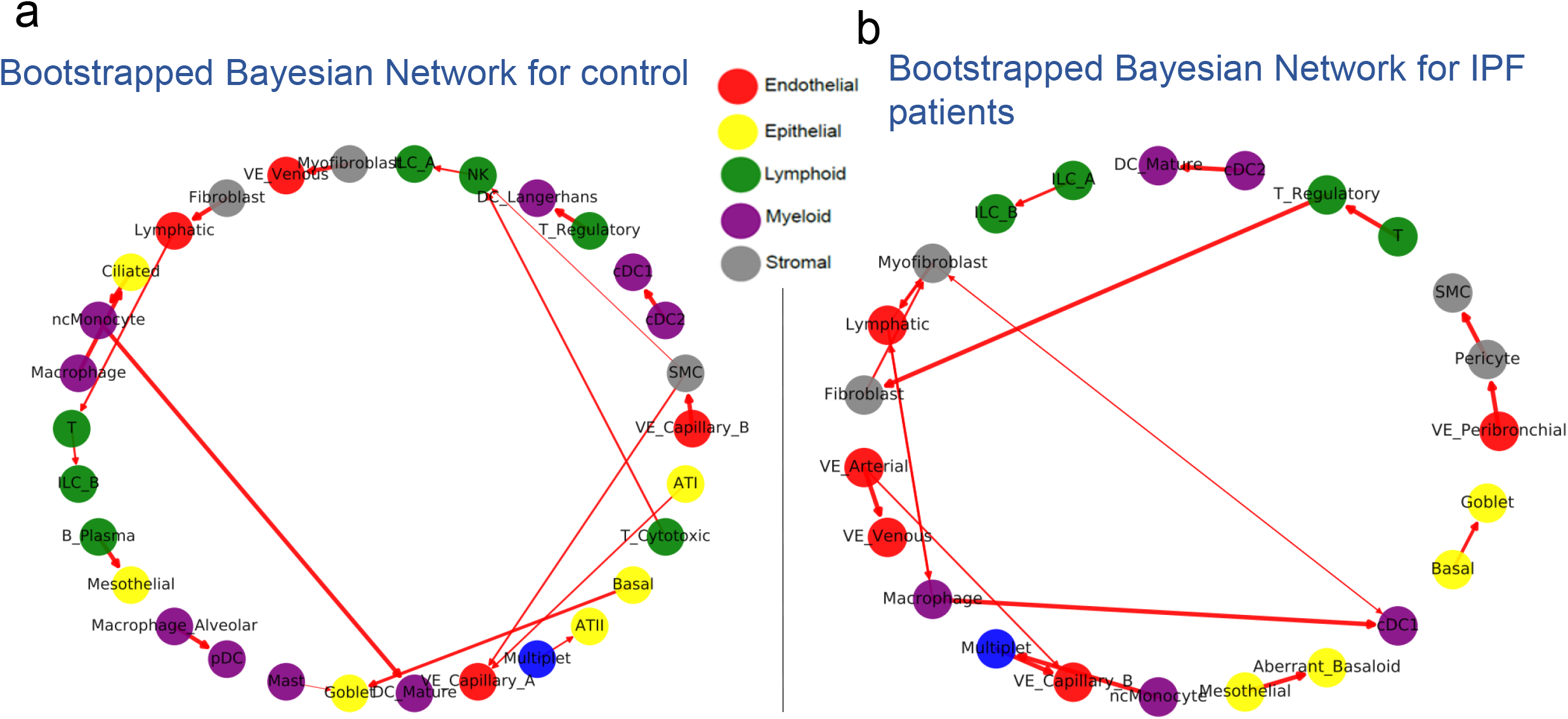
Bayesian Networks (BN) learned for lung cell types in healthy and IPF individual. (a) BN for controls (healthy individuals). (b) BN for IPF patients. Nodes represent specific cell types and are colored accordingly, edges represent directed interactions between the cell types. Edge width corresponds to its bootstrap score.

**Tab. 1.**
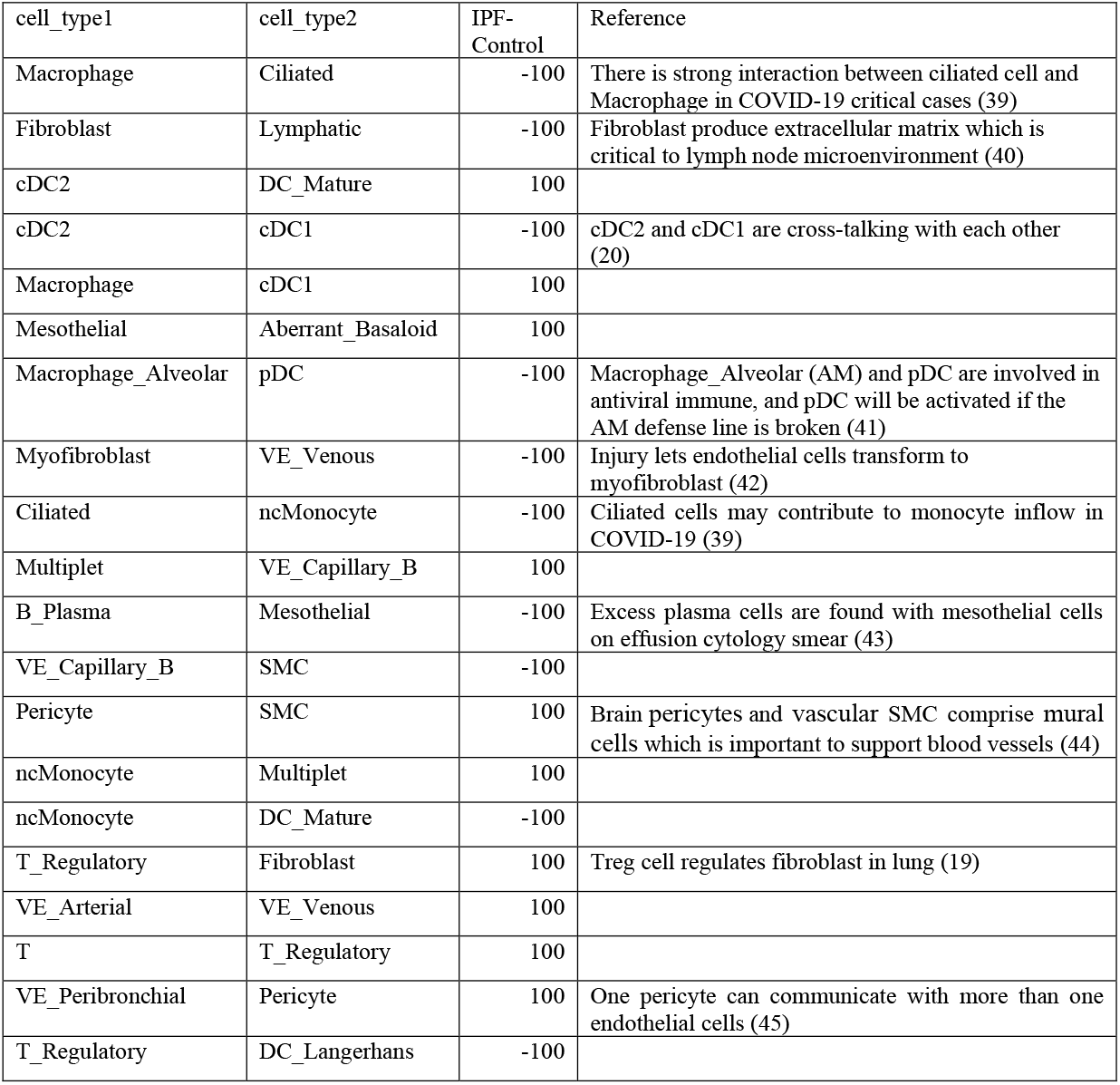
Top differential cell type interactions identified by CINS for the IPF dataset. The IPF-Control column lists the difference in the number of times the edge between the two cells was identified in 100 bootstrap runs for each of the two datasets. Negative values indicate that it was identified more for the Control whereas positive numbers mean that the interaction is more prevalent in IPF. For all listed edges the interaction was only identified in for one of the two datasets (score of 100 or −100).

### Inferring ligand-target interactions for significant cell type pairs

While the BNs discussed above identify pairs of cell types that likely interact in disease, the network does not show which genes and protein products participate in the interactions. To infer such gene-gene interactions across cells we developed a ligand-target regression (LTR) model. For cell type pairs identified in the BNs our LTR model uses a set of ligands in the first cell type to predict the expression values of their known targets in the second cell type. The LTR model uses the LASSO algorithm which enables the identification of a small set of key ligands predicted to participate in the interaction observed in the BN. We trained the model using a five-fold cross validation strategy. See **Methods** for details.

The LTR method was applied to all significant pairs identified by the BN. **Tab. S2** presents top scoring ligands for several cell type pairs. **Tab. S3** presents top scoring ligands for one cell type pair (Fibroblast -> Lymphatic cell). Several of the top LTR ligands are known to play an important role in the activated cell (Lymphatic cell). For example, the highest scoring ligand identified by LTR is “FGF2” which was identified as a critical gene for lymphangiogenesis (21). Another highly ranked ligand, “TGFB1”, can also accelerate lymphatic regeneration in wound repair (22). **Tab. S4** presents top ranked ligands for another pair (Treg cell -> Fibroblast), several of which have also been shown to participate in the interaction between these cell types. For example, fibroblast express IL13 receptor and may behave as an inflammatory cell if stimulated by IL-13 (23), and TGFB1-3 (including TGFB1 and TGFB2 in the table) are all involved in promoting collagen production in fibroblasts (24).

### Identified ligands are primarily involved in cell-cell interactions

To test if the predicted ligands are indeed impacting cell type-cell type interactions or mainly represent autocrine relationships we compared the activity of top predicted ligands within and between cell types. For this, we compared the performance of the LTR method for top edges to the performance of a similar method that only uses information from a single cell type. Specifically, if the BN predicted a significant interaction between cell types A -> B, we first trained LTR using the ligands of A and the targets of B (as we did above) and compared the performance to a LTR model which uses the ligands expressed in B to predict targets in B (autocrine model).

Results for the significant edges in the IPF and control datasets is presented in **Fig. 3A. Fig. 3B.** presents the results for the same pairs (so x axis is fixed based on the BN significance) but with the LTR trained using only the ligands of the second cell type. As can be seen, when using the ligand of the predicted interacting cell type LTR obtained a higher average correlation with a p-value of 0.034 (using the scipy function in Python for computing Pearson correlation p-values). In contrast, when using the same cell type for both ligands and targets the Pearson correlation is much lower (**Fig. 3B)**. We also evaluated the performance of the LTR method on the predicted cell type interactions by comparing the results we obtained with the real ligand-target interaction matrix to results obtained using a random ligand-target interaction matrix. We found that for most of the random assignments the resulting LASSO models contained only a Bias term with all coefficients set to 0 (**Fig. S3**). This indicates that expression of the ligands did not provide any useful information about the expression of the targets when using the random interaction matrix.

**Figure 3.**
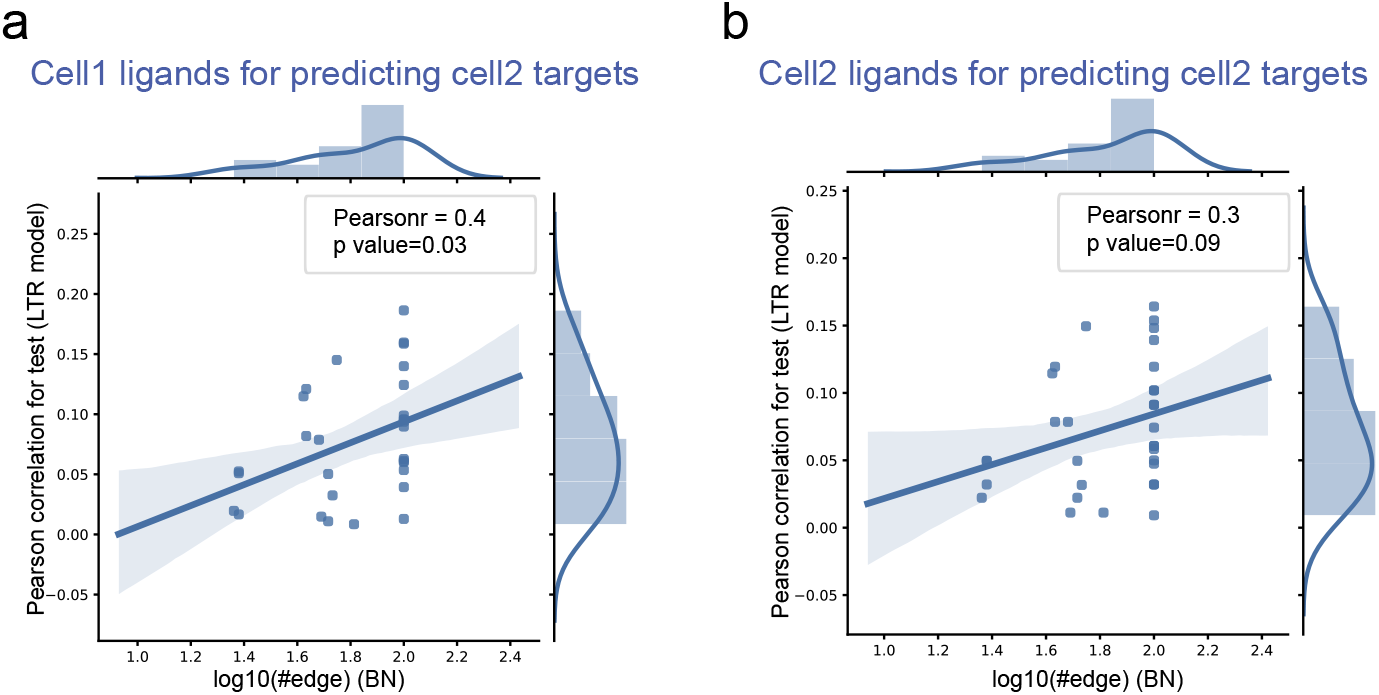
Interactions learned by the BN are more significant than interactions between cells of the same type. Comparison between the ability of the LTR model to predict target expression when learning the model using cells identified by the BN (a) and the same cell type (b). The x axis represents the bootstrapped edge count (significance) of the interaction in the BN for a cell type pair, and the y axis represents the LTR model performance (higher is better) for the same cell pair.

### Application to a scRNA-Seq dataset on lung aging

We next applied CINS to another, smaller, scRNA-Seq dataset which studied lung aging in mice (25). The dataset profiled lung cells in 15 mice, 8 young (three-month, 3M) and 7 old (24-month, 24M). The 14,813 cells profiled in this study were assigned to one of 34 cell types in the original paper. We again learned 100 bootstrapped BNs for the two conditions (young and old) and compared the resulting networks. We found 11 edges to be differentially present between the two conditions when using an edge threshold count of 20 (**Fig. 4** and **Tab. S5**). These included an edge between Capillary-endothelial-cell and Type 1-pneumocyte cells which are known to jointly form thin air-blood barriers used for gas exchange (26). Another pair was Ciliated and Club cells, of which the ratio is reported to alert significantly between young and old mouse lung (25). We next performed LTR analysis on the significant edges (**Fig. 5**). The top ranked ligand in Ciliated cells, TNF is known to regulate CC16 gene production, which plays a role in immunomodulatory activity in Club cells (27). Apoe, a ligand identified for the macrophage to goblet edge, is produced by macrophages to negatively modulate goblet cell hyperplasia (28).

**Figure 4.**
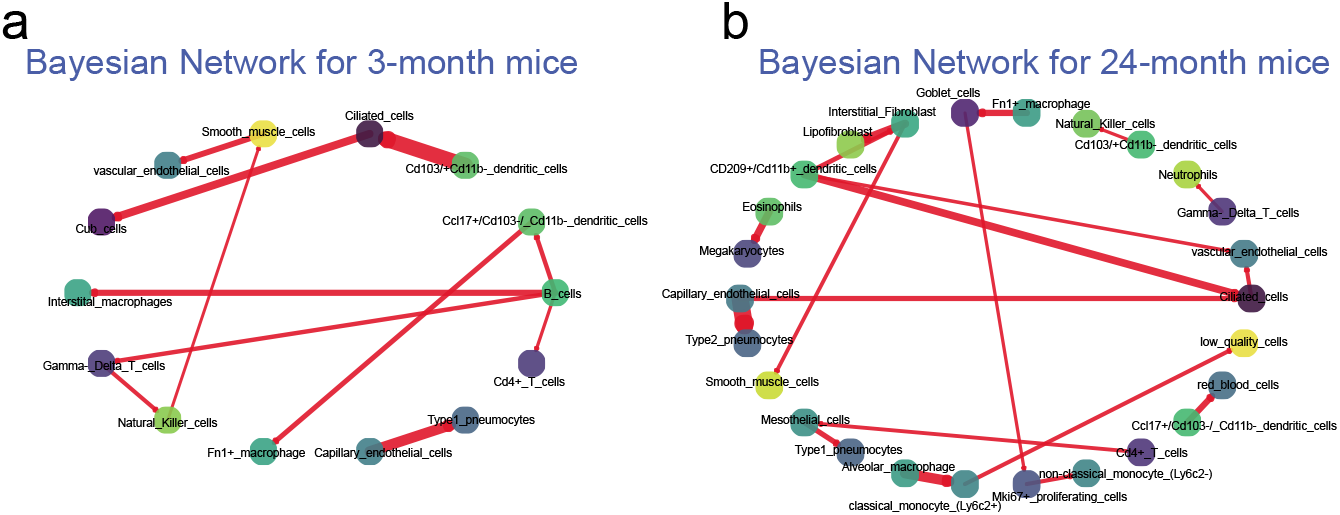
Aging Bayesian Networks. (a) BN for young mice. (b) BN for adult mice. Nodes and edges notations and colorings are similar to those used in **Fig. 2**.

**Figure 5.**
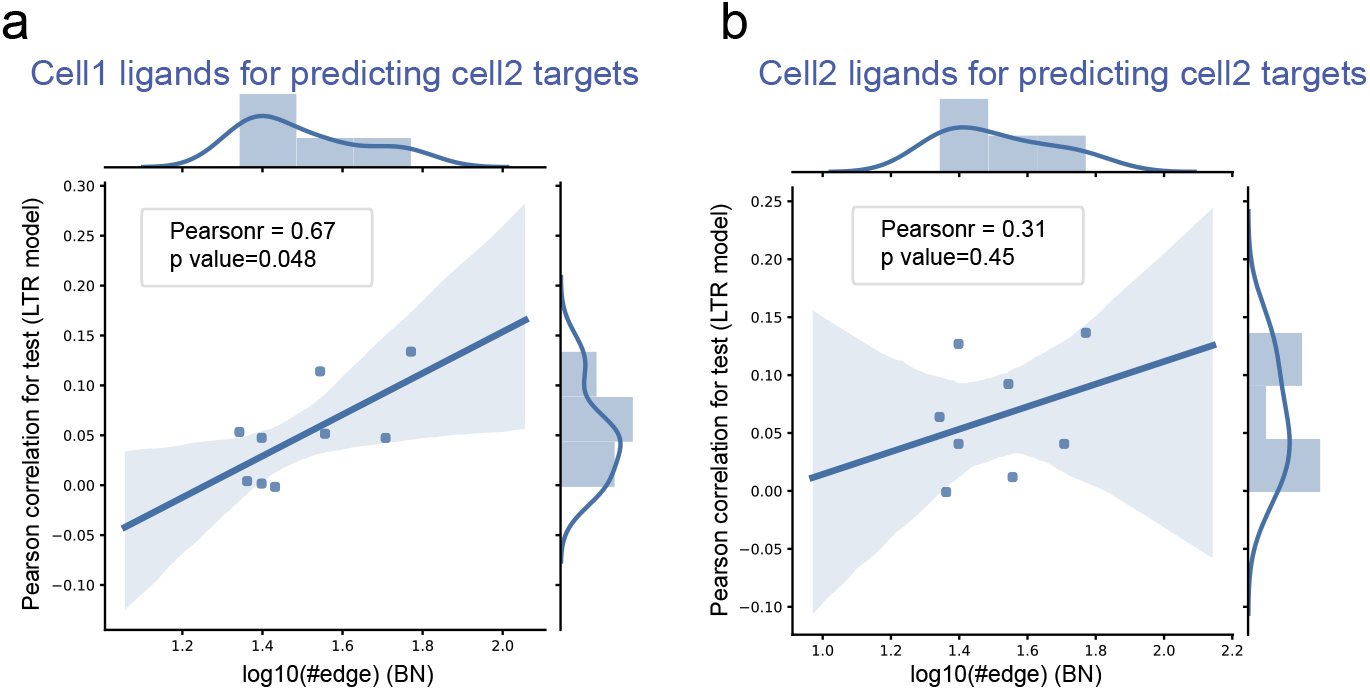
LTR comparison for the aging data. Comparison between the ability of the LTR model to predict target expression when learning the model using cells identified by the BN (a) and the same cell type (b).

As we did for the IPF study we compared the performance of the LTR method using ligands from the BN identified edges (A -> B) and ligands from the same cell type (B) to predict target expression for genes in B. We observed a Pearson correlation of 0.67 when using the ligands from vs. Pearson correlation of 0.31 when using the ligands from B. And it is noticed that when randomizing the interactions the LTR method again failed to identify any significant correlation between predicted and real expression for the targets (**Fig. S3**).

### Computational Validation of Significant Edges using a second aging mouse lung dataset

To test the predictions of the aging BN and to validate them using an independent cohort we next performed additional scRNA-Seq experiments on young and old mice to generate a pilot scRNA-Seq dataset on lung aging. For this, we profiled four young and four old mice of the *Fendrr-floxed* genotype recently generated in the Kaminski laboratory. We obtained 71,562 cells that were clustered, annotated, and assigned to 20 cell types that overlapped with the cell types assigned by Angelidis I et al. (25). We next used the combined data (from (26) and from our new experiments) to learn a joint BN. Several of the predicted interactions were further supported by our new data. Specifically, we found 19 cell type pairs for which the addition of our new data enhanced both the presence of the edge and the direction predicted when performing the bootstrap analysis. **Tab. S6** presents the top 10 enhanced pairs based on the overall bootstrap score (See **Tab. S9** for all enhanced pairs). For example, the interaction between Neutrophils and Gamma Delta T cell is enhanced from edge count of 40 to 61 and was reported by recent studies that neutrophils can suppress Gamma Delta T cell’s activation involved in the resolution of inflammation (29). And the interaction between B Cell and CD4+ T Cell is enhanced from −16 to −19 (being negative means that old lung has less), and is supported by other studies that B cell will activate CD4 T cells in human cutaneous leishmaniasis infection led by Viannia (30). In addition, we also found that T-cell-B-cell interactions were calculated to occur less often in older samples, which further validates the comparison between old and young mice (31).

We next focused on the top five predicted interactions in **Tab. S6** (all with an absolute enhanced bootstrap score larger than 15). Permutation analysis indicates that identifying such a large number of edges supported by both studies is significant (p-value = 0.05, **Methods** and **Fig. 6,** and see **Tab. S11** for result of other threshold values). We applied LTR to the cell type pairs in **Tab. S6** to find important ligand genes. **Tab. S7** presents the top predicted ligand genes. Several of these (red font) are supported by prior studies on the interaction between these cell types.

**Figure 6.**
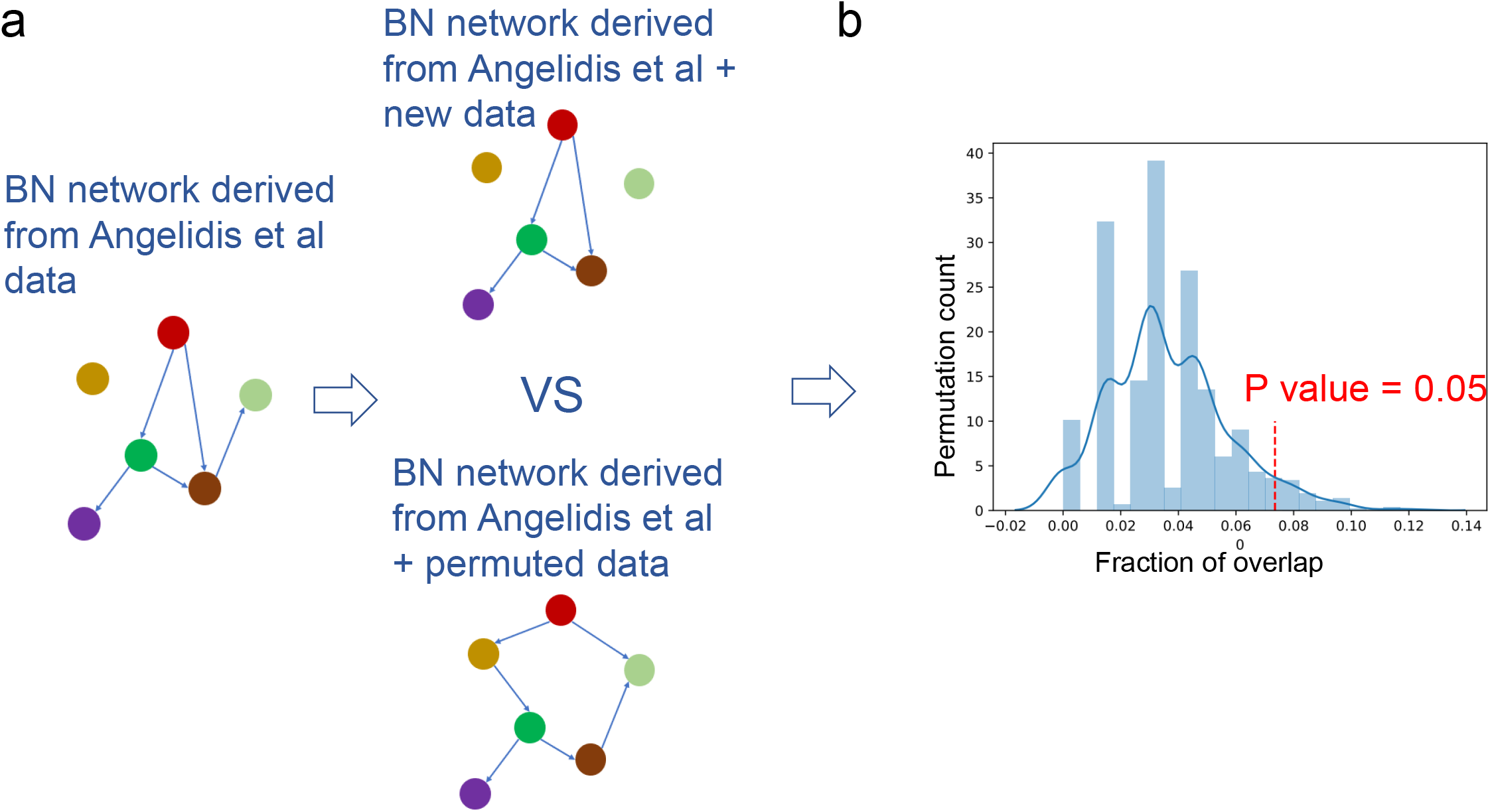
Permutation analysis highlights the agreement between the two aging networks. (a) Top – Learning combined networks using real data. Bottom – Learning combined networks using permutation analysis of cell type fractions in the new data. (b) Overlap in bootstrapped edges between the original and combined model when using the real data (red dashed line) and the permutation data (blue distribution).

## Discussion

To enable the study of cell type – cell type interactions using scRNA-Seq data we developed a method termed Cell Interaction Network Inference (CINS). CINS first learns a Bayesian network between cell types (BN) using repeated samples. Significant cell type pairs identified by the BN are further studied to infer the ligands that regulate these interactions. CINS is implemented in python and R and can be downloaded from www.github.com/CINS.

While CINS can be applied to any dataset with multiple samples, it is most appropriate for datasets containing case and control or multiple conditions. For such datasets CINS can infer not only the significant interactions within a condition but also those interactions that differ between the condition and that may partially explain the differences between the conditions studied.

We first applied CINS to study a case and control dataset profiling lung expression from IPF patients and controls. CINS identified several significant differences between the interactions observed for IPF patients and for healthy individuals. These include the interaction from Treg to Fibroblast cells which is supported by a recent study that found Treg can negatively regulate fibroblast activity (19), and the edge between cDC2 and cDC1 is also supported by recent work showing that cDC2 and cDC1 are cross-talking with each other (20).

For many of the identified significant interactions CINS was also able to identify key ligands involved in the interactions. For example, “FGF2” which was identified as a critical gene for lymphangiogenesis (21), and one more highly ranked ligand, “TGFB1”, can also accelerate lymphatic regeneration in wound repair (22).

We next applied CINS to a lung scRNA-Seq aging dataset and identified a number of significant pairs that differ between young and old mice. To validate predicted interactions we performed additional experiments in which we profiled scRNA-Seq expression in 4 additional young and old mice and then used the combined dataset to learn a joint network. As we showed, the network we learned identified a significant number of interactions that are supported by both datasets. These include the interactions between Neutrophils and Gamma Delta T cell (29), and between B Cell and CD4+ T Cell (30, 31) which are both supported by previous studies. CINS was again able to identify key ligands involved in these interactions, TNF, identified as the top ligand in the interaction between neutrophils and Gamma Delta T cells was previously identified as expressed in neutrophils (32) and as a regulator of immune cells Gamma Delta T cells (33), and TNFSF18 identified in interactions between CD4+ T cells and Vascular Endothelial Cells, was also previously reported to mediate the interactions between immune cells and endothelial cells (34).

While CINS can be successfully applied to several scRNA-Seq studies, it does have several limitations. First, it can only be applied if multiple samples are profiled since the BN part requires several repeated samples to compute relationships between cells. In addition, because BNs do not allow self edges interactions between cells of the same type cannot be identified by CINS. Finally, since it uses a bootstrap approach to infer significance it can miss important interactions if not enough samples and / or cells are available.

CINS is one of the first methods to enable the inference of cell type interactions in scRNA-Seq data from *repeated* samples. Given the growing popularity of this method, and its increased use in clinical studies which are currently less amenable to spatial transcriptomics techniques we believe that CINS provides a solution to an important problem that is not currently addressed.

## Methods

We developed a pipeline for modeling interactions between cells of different types from scRNA-Seq data. Our method first identifies cell types that are likely interacting and then tries to provide a mechanistic model to explain how such interactions are manifested at the molecular level.

### Datasets

We tested CINS using three scRNA-Seq datasets. The first compared gene expression in lungs of healthy and Idiopathic Pulmonary Fibrosis (IPF) with accession number of GSE136831 (15). This dataset contained 28 controls and 32 IPF patients with a total of 243,472 cells and the expression levels for 45,947 genes in each cell. We used the original annotations and included in the model all 39 cell types with at least 100 cells. The second dataset studied lung aging in mice with accession number of GSE124872 (25). This dataset contained 8 three-month-old mice and 7 24-month-old mice for which a total of 14,813 cells were profiled. For each cell the expression levels of 21,969 were provided. Each cell was assigned by the authors to one of 34 cell types. The third dataset was a new dataset in which we profiled single cell expression in four young (25 weeks) and four old (2x 103 weeks; 2x 120 weeks, Supporting Methods) *Fendrr-floxed* mouse lungs. This dataset contained a total of 71,562 cells with expression values for 45,947 genes. These cells were originally assigned to 37 cell types based on the expression of canonical cell type markers. To combine the two aging datasets we identified a joint subset of 20 cell types identified by both and only used cells assigned to on these cell types in our combined BN analysis.

Information about ligands and their targets were obtained from a recent paper (14) which provided targets for 688 ligands.

### Single-cell sequencing of *Fendrr-floxed* Mice

Animal procedures had been approved by the Institutional Animal Care and Use Committee (IACUC). We created a floxed allele of *Fendrr* via two-guide, two-oligo CRISPR/Cas mediated cleavage and recombination essentially as described in Yang et al. (35). A generated mouse which had the expected conditional allele was bred with C57BL/6J mice to establish the colony and to sort the floxed allele from any other possible mutant alleles. Three female and five male mice in two age groups (young: 23 weeks, old: ranging from 103 to 120 weeks; four mice per group) were euthanized, and lungs were harvested and minced in small pieces with a scalpel. Lung pieces were dissociated using the enzyme Liberase TL (Roche).

Single RNA molecules of single cells were barcoded using the 10× chromium single-cell technology according to the manufacturer’s instructions (Single Cell 3’ Reagent Kits v2, 10× Genomics, USA). Barcodes were used to assign reads to cell and quality control was performed to remove low quality cells (Supporting Methods). Generated sequencing data is available at GEO accession number GSE165638. A modified version of the standard Seurat pipeline was employed to normalize, cluster and annotate the raw counts single-cell expression data for downstream analysis (36). Briefly, the percent of mitochondrially-expressed genes was calculated for each individual cellbarcode using the *PercentageFeatureSet* function. Next, unique molecular identifier (UMI) counts were log normalized with a scale factor of 10,000 UMIs per cell and then natural log transformed using a pseudocount of one. Following log normalization, the top 3500 variable genes within the dataset were determined using Seurat’s implementation of the *FindVariableFeatures* function with the “vst” parameter. Next, the gene-level scaling of the data was performed using the *ScaleData* function. Each feature was centered to have a mean of zero and scaled by the standard deviation of each feature. The percent of mitochondrially-expressed genes captured within each cell were regressed out during scaling by using the “vars.to.regress” parameter. To reduce the dimensionality of the dataset and to identify genes contributing the most variability to the underlying manifold of the dataset, Principal Component Analysis (PCA) was performed using the scaled data and the 3500 variable genes calculated determined for the dataset. Following exploration of the PCs (Supporting Methods), the first 75 PCs were selected for clustering and UMAP projection. The quality of subject and age representation within each cluster was assessed prior to cell type annotation to note any subject- or age-specific biases.

### Cell type assignment of *Fendrr-floxed* mice

To assign a specific cellular identity to each cluster, differentially expressed markers were determined and assessed within the context of canonical marker genes. Briefly, a differential gene expression test using Wilcoxon Rank Sum test was performed that compared the gene expression within a specific cluster to expression within all cells outside of that cluster. The resulting list of cluster-specific marker genes was assessed and cell types were ascribed based on expression of canonical marker genes. Clusters displaying canonical markers for multiple cell types were flagged as multiplets and were omitted from downstream analysis.

### Cell type quantification and discretization

We use the cell type annotations provided by each of the study. To use Bayesian network to learn relationships between cell type we first discretize the proportion of each cell type in each sample. Discretization is cell type specific (i.e. different cell type will be assigned different values for the same proportion quantity) and is learned using an unsupervised method based on Gaussian Mixture Model (GMM) with two components. Specifically, let 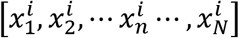 be the fraction (percentage) of the *ith* cell type in the N samples. We learn a two components GMM for these values and then assign each value to the class with the higher likelihood for this value. The target function of the GMM aims to maximize the log likelihood:

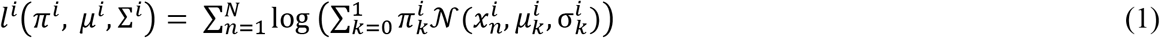

Where *N* represents gaussian distribution and 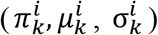 represent proportion, mean and standard deviation parameters for the *kth* component of the *ith* cell type.

Following convergence, each proportion value 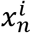 is assigned to one of the two classes. We assign labels to the two classes such that the component with lower mean parameter is assigned a value of 0 and the second is assigned a value of 1. This leads to a learned cell type specific cutoff such that all samples with a value less than that cutoff are assigned to 0 and all those above are assigned to 1. However, the number of 0’s and 1’s is not pre-determined and may be highly skewed in either direction based on the distribution of the fractions. See **Fig. S1** for examples of assignments. To learn GMMs we used the Python package “sklearn” with a maximum iteration number of 500 and a convergence threshold of 10**-4.

### Learning a cell type Bayesian network

We use the discretized cell type values to learn a cell type Bayesian network. Bayesian network is a probabilistic graphical model that uses directed acyclic graph to represent joint probability distributions. The absence of an edge can indicate independence and / or conditional independence. Bayesian networks are parameterized as <*G, P*> where G = <*V, E*> is a directed acyclic graph with *V* as variables and E as directed edges, and P is conditional probability for each node:

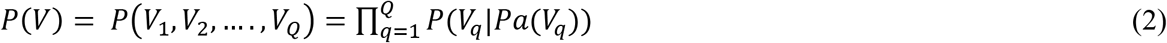

Where *Pa*(*V_q_*) is parent node set of *V_q_* according to *G*.

To learn a Bayesian network using the discretized cell type proportion data. we iterate between network learning and parameter estimation. We initialize the network using the *Hiton Parents and Children* strategy which is based on marginal association among variables (37). Next we iterate a search strategy, that uses penalized Hill-Climbing to add, flip or remove edges based on the Bayesian Information Criterion (BIC) score:

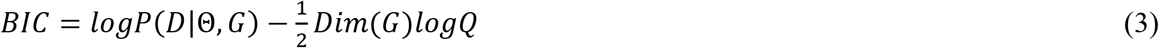

For this, we used the “rsmax2” function from the R library “bnlearn”, which implements the iterative Penalized Maximization algorithm to construct a Bayesian network.

To obtain confidence values for edges (predicted interactions) in the network we followed previous learning methods that utilized a bootstrap strategy (16, 17). For each iteration of the bootstrap we first randomly sample 80% of all single cells in the dataset. Next, we used these cells to determine cell type frequencies in each sample and to perform the discretization and network learning as described above. This step is repeated 100 times, and for which we counted the presence of all directed edges. While the direction of an edge in a Bayesian network does not always imply casual interactions (38), we observed that significant edges were also very consistent in their direction (**Tab. S1**).

### Ligand-Target Regression (LTR) Model

The bootstrapping method presented above provides a small set of significant interactions between some of the cell types in the dataset. To obtain a mechanistic explanation for these interactions, and to identify the interacting genes between the two cell types we focused on ligand-target interactions. Specifically, for a directed edge between cell A and B we learned a regression model to explore the underlying gene interactions between A and B. Our assumption is that if these two cell types indeed interact, then ligand gene expression data in cell A should be able to explain some of the expression changes observed in cell type B. To identify the set of ligands in A predicted to activate or repress genes in B we optimized the following model:

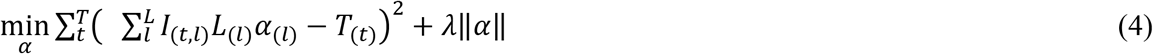

Where *I* represents an input (known) ligand-target interaction matrix (14), *L* is a vector of log values for the ligand expression fold change levels in cell type A, *α* represents the (unobserved) ligand activation vector, *T* represents log value of gene expression fold change levels for target genes in cell type B and *λ* is a penalty parameter. Here we used a *L1* penalty which usually leads to the selection of relatively few non zero values (corresponding to relatively few activated ligands in cell type A).

Setting *A*_(*t,l*)_ = *I*_(*t,l*)_ *L*_(*l*)_, transforms the optimization problem to

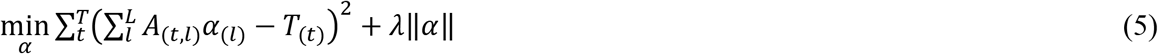

Which is a standard least absolute shrinkage and selection operator (LASSO) model. For this, we used the “LASSOCV” function from the Python library “scikit-learn”, which implements the LASSO cross validation.

### Training and Test for Ligand-Target Regression (LTR) Model

We used a five-fold cross validation to train and test the optimization model: We split the dataset into three parts, training, validation and test. We first use the validation set to select the optimal penalty term *λ* and then retrain the using the entire data and the selected *λ* to obtain the model used for the fold test data. Evaluation of predicted values is based on the average Pearson correlation between the predicted and actual expression changes for each fold. Following testing we use the average product between the log fold change and coefficient value *α* in the five-fold training models to rank the list of active ligands.

### Joint plots of Bayesian Network and LTR model scores for cell type pairs

To jointly plot the Bayesian network bootstrap score and the Pearson correlation regression score for each cell type pair, we first converted the edge count to log value. For the Pearson correlation we used the average correlation for the five-fold results. For both IPF lung data and lung aging data, cell pairs with edge count smaller than 20 are removed (See **Tab. S10** for details). Note that for some of the pairs we tried to model using LASSO the learning terminated with coefficients of 0 for all ligands (this happened for all runs of the random interaction matrix as we mention in Results and to a few of the CV runs of the cell-cell and intra-cell models). In such cases these models were removed from the correlation analysis.

## Acknowledgements

Work partially funded by the National Institutes of Health (NIH) (https://www.nih.gov) [grants 1R01GM122096 and OT2OD026682 to Z.B.J. and 1R01HL127349 to N.K.].

## Author contributions

Y. Y., M.R. and Z.B.-J. designed research; Y.Y., C.C.J., T.S.A., J.S., K.S., N.X., M.R., N.K., and Z. B.-J. performed research; C.C.J., T.S.A., J.S., K.S., N.X., M.R., N.K. generated data; Y.Y. analyzed data; and Y.Y., C.C.J., and Z.B.-J. wrote the paper. All authors read and approved the paper.

## Data Availability

All scripts, and instruction required to run CINS pipeline in Python and R can be found in our support website, https://github.com/xiaoyeye/CINS. Generated sequencing data is available at GEO accession number GSE165638. All other public data can be found following the pipelines in Methods.

## References

1. Deng Y, Bao F, Dai Q, Wu LF, & Altschuler SJ (2019) Scalable analysis of cell-type composition from single-cell transcriptomics using deep recurrent learning. Nat Methods 16(4):311–314.

2. Alavi A, Ruffalo M, Parvangada A, Huang Z, & Bar-Joseph Z (2018) A web server for comparative analysis of single-cell RNA-seq data. Nat Commun 9(1):4768.

3. Kumar MP, et al. (2018) Analysis of Single-Cell RNA-Seq Identifies Cell-Cell Communication Associated with Tumor Characteristics. Cell Rep 25(6):1458–1468 e1454.

4. Han X, et al. (2018) Mapping human pluripotent stem cell differentiation pathways using high throughput single-cell RNA-sequencing. Genome Biol 19(1):47.

5. Moncada R, et al. (2020) Integrating microarray-based spatial transcriptomics and single-cell RNA-seq reveals tissue architecture in pancreatic ductal adenocarcinomas. Nat Biotechnol 38(3):333–342.

6. Codeluppi S, et al. (2018) Spatial organization of the somatosensory cortex revealed by osmFISH. Nature methods 15(11):932.

7. Lee JH, et al. (2014) Highly multiplexed subcellular RNA sequencing in situ. Science 343(6177):1360–1363.

8. Moffitt JR, et al. (2018) Molecular, spatial, and functional single-cell profiling of the hypothalamic preoptic region. Science 362(6416):eaau5324.

9. Ståhl PL, et al. (2016) Visualization and analysis of gene expression in tissue sections by spatial transcriptomics. Science 353(6294):78–82.

10. Eng C-HL, et al. (2019) Transcriptome-scale super-resolved imaging in tissues by RNA seqFISH+. Nature 568(7751):235.

11. Xia C, Fan J, Emanuel G, Hao J, & Zhuang X (2019) Spatial transcriptome profiling by MERFISH reveals subcellular RNA compartmentalization and cell cycle-dependent gene expression. Proceedings of the National Academy of Sciences 116(39):19490–19499.

12. Dries R, et al. (2019) Giotto, a pipeline for integrative analysis and visualization of single-cell spatial transcriptomic data. bioRxiv.

13. Yuan Y & Bar-Joseph Z (2019) GCNG: Graph convolutional networks for inferring cell-cell interactions. bioRxiv:2019.2012.2023.887133.

14. Browaeys R, Saelens W, & Saeys Y (2019) NicheNet: modeling intercellular communication by linking ligands to target genes. Nature Methods:1–4.

15. Adams TS, et al. (2020) Single-cell RNA-seq reveals ectopic and aberrant lung-resident cell populations in idiopathic pulmonary fibrosis. Sci Adv 6(28):eaba1983.

16. Lugo-Martinez J, Ruiz-Perez D, Narasimhan G, & Bar-Joseph Z (2019) Dynamic interaction network inference from longitudinal microbiome data. Microbiome 7(1):54.

17. Imoto S, et al. (2002) Bootstrap analysis of gene networks based on Bayesian networks and nonparametric regression. Genome Informatics 13:369–370.

18. Kia’i N & Bajaj T (2020) Histology, Respiratory Epithelium. StatPearls, Treasure Island (FL)).

19. Qiu H, et al. (2019) The Role of Regulatory T Cells in Pulmonary Arterial Hypertension. J Am Heart Assoc 8(23):e014201.

20. Noubade R, Majri-Morrison S, & Tarbell KV (2019) Beyond cDC1: Emerging Roles of DC Crosstalk in Cancer Immunity. Front Immunol 10:1014.

21. Platonova N, et al. (2013) Evidence for the interaction of fibroblast growth factor-2 with the lymphatic endothelial cell marker LYVE-1. Blood, The Journal of the American Society of Hematology 121(7):1229–1237.

22. Avraham T, et al. (2010) Blockade of transforming growth factor-beta1 accelerates lymphatic regeneration during wound repair. Am J Pathol 177(6):3202–3214.

23. Doucet C, et al. (1998) Interleukin (IL) 4 and IL-13 act on human lung fibroblasts. Implication in asthma. J Clin Invest 101(10):2129–2139.

24. Coker RK, et al. (1997) Transforming growth factors-beta 1, -beta 2, and -beta 3 stimulate fibroblast procollagen production in vitro but are differentially expressed during bleomycin-induced lung fibrosis. Am J Pathol 150(3):981–991.

25. Angelidis I, et al. (2019) An atlas of the aging lung mapped by single cell transcriptomics and deep tissue proteomics. Nature communications 10(1):1–17.

26. Kara Rogers Senior Editor BS (2010) The Respiratory System (Britannica Educational Pub.).

27. Bergamaschi E, Canu IG, Prina-Mello A, & Magrini A (2017) Biomonitoring. Adverse effects of engineered nanomaterials, (Elsevier), pp 125–158.

28. Acton QA (2013) Asthma: New Insights for the Healthcare Professional: 2013 Edition (ScholarlyEditions).

29. Sabbione F, et al. (2014) Neutrophils suppress gammadelta T-cell function. Eur J Immunol 44(3):819–830.

30. Rodriguez-Pinto D, Saravia NG, & McMahon-Pratt D (2014) CD4 T cell activation by B cells in human Leishmania (Viannia) infection. BMC Infect Dis 14:108.

31. Salam N, et al. (2013) T cell ageing: effects of age on development, survival & function. Indian J Med Res 138(5):595–608.

32. Grivennikov SI, et al. (2005) Distinct and nonredundant in vivo functions of TNF produced by t cells and macrophages/neutrophils: protective and deleterious effects. Immunity 22(1):93–104.

33. Lahn M, et al. (1998) Early preferential stimulation of gamma delta T cells by TNF-alpha. J Immunol 160(11):5221–5230.

34. Anonymous (2020) TNFSF18 TNF superfamily member 18 [Homo sapiens (human)].

35. Yang H, Wang H, & Jaenisch R (2014) Generating genetically modified mice using CRISPR/Cas-mediated genome engineering. Nat Protoc 9(8):1956–1968.

36. Stuart T, et al. (2019) Comprehensive Integration of Single-Cell Data. Cell 177(7):1888–1902 e1821.

37. Aliferis CF, Statnikov A, Tsamardinos I, Mani S, & Koutsoukos XD (2010) Local causal and markov blanket induction for causal discovery and feature selection for classification part i: Algorithms and empirical evaluation. Journal of Machine Learning Research 11(Jan):171–234.

38. Nadkarni S & Shenoy PP (2001) A Bayesian network approach to making inferences in causal maps. European Journal of Operational Research 128(3):479–498.

39. Chua RL, et al. (2020) COVID-19 severity correlates with airway epithelium-immune cell interactions identified by single-cell analysis. Nat Biotechnol 38(8):970–979.

40. Morgado FN, da Silva AVA, & Porrozzi R (2020) Infectious Diseases and the Lymphoid Extracellular Matrix Remodeling: A Focus on Conduit System. Cells 9(3).

41. Kumagai Y, et al. (2007) Alveolar macrophages are the primary interferon-alpha producer in pulmonary infection with RNA viruses. Immunity 27(2):240–252.

42. Chaudhuri V & Karasek MA (2006) Mechanisms of microvascular wound repair II. Injury induces transformation of endothelial cells into myofibroblasts and the synthesis of matrix proteins. In Vitro Cell Dev Biol Anim 42(10):314–319.

43. Gochhait D, Dey P, & Verma N (2016) Cytology of plasma cell rich effusion in cases of plasma cell neoplasm. J Cytol 33(3):150–153.

44. Smyth LCD, et al. (2018) Markers for human brain pericytes and smooth muscle cells. J Chem Neuroanat 92:48–60.

45. Sweeney M & Foldes G (2018) It Takes Two: Endothelial-Perivascular Cell Cross-Talk in Vascular Development and Disease. Front Cardiovasc Med 5:154.

